# Modeling Synthetic Audience Reactions to Narratives: A Generative GSR Model to Predict Autonomic Alignment during Film Viewing

**DOI:** 10.64898/2026.04.13.716763

**Authors:** Benjamin Bartling, Hee Jung Cho, Yuetong Du, Ralf Schmälzle

## Abstract

While media consumption can be a solitary act, it produces a shared, socially coordinated experience where audiences’ bodies align in response to shared narrative events that are often social-affective in nature. Despite this recognition, traditional descriptive models of Galvanic skin response (GSR) have existed for decades, yet the socially coordinated aspect remains to be fully reflected in physiological models with the field of communication often treating the underlying generators of autonomic activity as a black box. To bridge this gap, we introduce a computational framework that models the underlying neural driver and its convolution to sweat gland physiology to explain how narrative events translate into measurable conductance. By leveraging multimodal AI models to “interpret” the social-cognitive content of a film, we generated a predictor timeline for a synthetic audience comprised of digital agents (i.e. artificial body systems responding to the film events with GSR responses). We then test this computational audience model by comparing its predictions against an empirical dataset collected as audience members (*N* = *96*) processed the same stimulus, finding that AI-identified social triggers, like moments of comedic violence or shared emotional shifts, significantly predict the GSR time-course of audience engagement. In sum, this paper moves beyond simple and often retrospective labels like “arousal” to offer a computational account of how shared social narratives grip the human nervous system. We provide a scalable and expandable framework and a set of tools to predict media impact and understanding the psychophysiological basis of media.

---

The study of human responses to dynamic media has long relied on Galvanic Skin Response (GSR) as a proxy for emotional arousal (Lang, 2009). Despite this reliance, a significant methodological rift remains: We lack a generative model that can simulate realistic GSR traces based on specific media events. While such approaches exist for fMRI and eye-tracking (Zeidman et al., 2025; Itti and Koch, 2000), GSR modeling has remained more descriptive in nature, and the few model-based approaches have utilized conditioning paradigms (Benedek and Kaernbach, 2010a; Bach et al., 2010). Advancing the field requires shifting toward dynamic, mechanistic models capable of generating realistic, stimulus-driven GSR time-series from continuous narratives that are key to social-affective research. Rather than merely describing when arousal occurs, the next step is to model how autonomic responses dynamically unfold in response to structured media input and how these dynamics support social cognitive processes by aligning audience physiological responses over time.

This paper introduces a computational framework for simulating GSR responses to the Pixar short film Partly Cloudy. This Pixar short film provides an ideal stimulus due to its structured tension-resolution cycles, emotionally salient inflection points, and minimal dialogue allowing for precise extraction of stimulus dynamics. By modeling the underlying neural driver and its convolution through sweat gland physiology, we aim to explain how salient narrative events translate into measurable conductance. We utilize an iterative process that allows for the fine tuning of parameters, allowing us to study how this happens in single individuals and audiences at large, whose response similarity is examined via inter-subject correlation. In the following, we first introduce the role of GSR in communication, social cognition, and media psychophysiology research. We also discuss related model-based approaches in other domains of psychophysiology and neuroscience. We then introduce the current study and compare simulated and real data.

## Literature Review

### GSR in Communication and Media Psychology

In the field of communication and media psychophysiology, skin conductance measurements1 serve as an index of emotional arousal (Potter and Bolls, 2012). Unlike heart rate, which is subject to both sympathetic and parasympathetic influences, skin conductance is uniquely innervated by the sympathetic nervous system, providing a relatively unobstructed measure of the receiver’s level of arousal in response to media (Boucsein, 2013; Lang, 2009), particularly during moments of high emotional or narrative intensity. GSR measures can trace the fluctuations in audience arousal during film viewing. Electrodermal activity tracks both immediate reactions to stimuli as well as sustained psychological states. Stimulus-driven immediate responses are captured by phasic measurements, known as the skin conductance responses (SCRs) (Bailey, 2017; Stern et al., 2001), whereas slower tonic changes are generally referred to as skin conductance level (SCL). Prior work has measured skin conductance to examine synchrony in arousal during ongoing media reception (e.g., Golland et al., 2014; Han et al., 2022).

### The Physiology of Skin Conductance and Lack of a Generative Model

Electrodermal activity is the only psychophysiological measure entirely governed by the sympathetic nervous system (Boucsein, 2013), as eccrine sweat glands respond exclusively to sympathetic input. These glands are primarily located on the palms and soles and are innervated by sudomotor nerves. In brief, when the brain detects a salient or semantically arousing stimulus, central autonomic networks, including regions such as the amygdala, anterior cingulate cortex, and hypothalamus (Boucsein, 2013; Critchley et al., 2000) prompt the activation of sympathetic sudomotor pathways. This results in sweat secretion, altering the skin’s electrical conductance. Because the signal results from sympathetic activation, electrodermal activity provides a window into autonomic arousal during media exposure.

Despite this relatively straightforward physiological chain, the field of communication and media psychophysiology has suffered from the lack of a formal, explicit generative model. While other domains have moved toward forward modeling (e.g. fMRI, eye-tracking, see below), where one predicts the observed signal by simulating the underlying biological process, GSR research remains largely descriptive. In GSR analysis, peaks are simply counted and reverse correlated with content after the fact. While sophisticated toolkits like Ledalab and PSPM/SCRalyze exist (Benedek and Kaernbach, 2010b; Bach, 2014b), they are primarily used for deconvolution, which is working backwards from a noisy signal to find the underlying spikes of engagement. Furthermore, these models were mostly developed for discrete, isolated stimuli in controlled experimental designs, such as fear conditioning. Therefore, when these methods have been used for deconvolution or the corresponding forward convolution, this has been done with simple stimuli and tasks very different from those we find in social-cognitive media psychology, where the stimuli are continuous, events may overlap or succeed each other rapidly, and eliciting stimuli are more complex and meaning-laden.

As long as we do not model the forward process, from narrative event to sudomotor burst to sweat gland response, we are prone to treat the GSR as a “black box”. Perhaps as a result of this void, there is a tendency to assume a rather simple one-to-one relationship between the measured “conductance” and the inferred “arousal” that supposedly generated the arousal (so-called reverse inference; Poldrack, 2011). Building a generative model that can simulate continuous GSR traces in a forward manner (from stimulus to brain activity to sudomotor nerve activity to measured skin conductance), allows us to overcome this theoretical bottleneck. Rather than attributing GSR to arousal, a rather broad and primitive construct, we can begin with the stimulus itself via content-analysis, one of our field’s backbone methods. For instance, there could be a surprising auditory event like a clap that would elicit a response, but also a salient emotional event, like two friends hugging; both would likely evoke a skin conductance response.

In this vein, modeling comes with key theoretical and methodological benefits (see Epstein, 2008): Perhaps most importantly, introducing a forward logic that goes from identified stimulus content to the measured physiological response increases the conceptual precision and theoretical clarity of media psychophysiological research (DeAndrea and Holbert, 2017). Moreover, we can simulate experimental effects, test the sensitivity of our analysis pipelines, estimate noise ceilings, gauge power, and so forth – all before even a single participant is ever recruited. We can also model additional parameters beyond content-based elicitors, such as various individual differences like variation in the (synthetic) audiences’ level of attentiveness, their physiological reactiveness, and various other parameters increasing conceptual clarity and theoretical precision. Collectively, computational modeling of SCR during narrative media represents an obvious and promising next step in the theoretical development of media psychophysiology.

### Related Frameworks in Psychophysiology: Lessons from fMRI and Eye-Tracking

The transition from purely descriptive to generative modeling follows a well-established trajectory in other domains of neuroscience and psychophysiology, where researchers have long sought to bridge the gap between an unobserved generator and its measurable output (Cacioppo et al., 2007).

Relationship to fMRI: The SPM framework and BOLD-response Convolution: Perhaps the most direct parallel to our GSR framework is the analysis of blood-oxygen-level-dependent (BOLD) signals in fMRI (Zeidman et al., 2025). In the widely used Statistical Parametric Mapping (SPM) framework, the timing of experimental stimuli is represented as a series of events in a design matrix. These events (spikes) are then convolved with a hemodynamic response function (HRF), which is a mathematical model of the delayed and dispersed way blood flow changes following neural activity. Our use of the Bateman function (see Methods) to model the sweat gland impulse response (see Methods) mirrors this HRF approach, treating the peripheral autonomic system as a filter that transforms discrete sympathetic bursts into recordable continuous signals.

Relationship to Eye-Tracking and Saliency Models: In the domain of visual attention, generative models have successfully predicted where people look before the data is ever collected (Itti and Koch, 2001; Berlot et al., 2026). Saliency models use computational algorithms to analyze the formal features of a stimulus, such as contrast, color, and motion, to generate a prediction map of human gaze (Walter and Bex, 2022; Walter et al., 2024). Our framework extends this logic to the autonomic layer, using narrative intensity as a saliency feature that elicits activity in the sympathetic nervous system. In doing so, we shift from predicting spatial patterns of attention to modeling the temporal dynamics of arousal during social cognition, thereby capturing how structured media input gives rise to shared physiological responses across viewers. This extension positions autonomic alignment as a form of socially coordinated processing, analogous to how visual saliency organizes attention in space. By treating narrative intensity as a form of affective saliency, this approach links low-level physiological responses to higher-order social-cognitive phenomena, suggesting that shared autonomic dynamics may serve as a mechanistic basis for collective experience during media consumption.

Related work in Skin Conductance Research: While the application of generative, continuous models to dynamic media is relatively new, the field of psychophysiology has a rich history of model-based SCR analysis. Traditional approaches, such as Ledalab and SCRalyze (now PSPM; Benedek and Kaernbach, 2010a; Bach, 2014a), paved the way by introducing model-based analyses for skin conductance data. These tools typically rely on de-convolution, the mathematical inverse of the process we employ here (forward convolution), to overcome the smearing effect of the sweat glands and in an effort to “recover” the underlying nerve activity that triggers the response.

These models have predominantly focused on discrete, well-separated stimuli in controlled experimental paradigms (Benedek and Kaernbach, 2010a; Bach, 2014a). For decades, dynamic media were considered too complex for such rigorous modeling due to overlapping responses of continuous narrative signals (Spiers and Maguire, 2007). Our work leverages the principles established in these earlier tools but applies them in a forward direction to simulate the complex, overlapping dynamics of an audience engaged with a naturalistic stimulus. Doing so is made possible by recent advances in AI research, where modern multimodal models are able to “view” content and generate predictors.

## The Current Study

To summarize, descriptive models of Galvanic Skin Response (GSR) have existed for decades. However, the field lacks a generative framework capable of predicting continuous physiological traces from exposure to temporally unfolding media messages. Existing approaches often rely on event locked analyses and post hoc signal detection, leaving the mechanistic transformation of narrative structure to autonomic dynamics in the black box of the brain.

The study reported below is directed at answering the following questions: Does our current theoretical knowledge about GSR warrant a computational framework to generate synthetic GSR data? Can we use current AI models to identify events in a complex, social movie for which GSR responses can be simulated? If so, can we simulate whole psychophysiological audiences and will synthetic data from a simulated GSR audience correlated with real data? What audience characteristics beyond the basic GSR physiology should be modeled (e.g. attentional fluctuations, variability in emotional reactivity, social comprehension)?

To address these issues, we introduce a computational framework that models the underlying neural driver and its convolution through sweat gland physiology to explain how narrative events during an animated movie translate into measurable conductance. This simulated agent allows us to not only evaluate individual responses but to instantiate a synthetic audience. By methodically parameterizing variability in sensitivity, latency, recovery rate, and noise structure, multiple simulate agents can be examined collectively. Thus, this approach enables the modeling of predicted autonomic alignment across viewers and to compute ISC directly from simulated data.

Then, we validate this model by comparing the synthetic signals and predicted alignment patterns against an empirical dataset collected as part of a large-scale study. This dual approach of simulation and empirical comparison allows us to quantify the validity of our simulation and test its ability to generate the hierarchical architecture of human autonomic responses from momentary stimulus locked signals, individual level variability, and to collective audience synchronization.

## Methods

This section is organized into the following subsections that bridge computational modeling with empirical validation. First, we detail the GSR Simulation Framework, which uses a biophysical approach to synthesize skin conductance signals. Second, we describe the Movie Stimulus and how it was analyzed using multimodal AI and a dedicated prompt to generate the time-line eliciting events that are then fed into the model to simulate what GSR data would look like. Then, we describe how we used the data from the movie annotation stage as input to the GSR simulation framework to generate synthetic audience data. Finally, we provide a description of the real-world GSR dataset, which served as the comparison standard to evaluate the quality of the model-based simulation.

### Simulating GSR Responses

The simulation framework was implemented in Python using a biophysical, forward-modeling approach to transform discrete narrative events into continuous, population-level physiological signals. The modeling process follows three primary stages: individual agent parameterization, neural driver generation, and physiological convolution.

Bateman Impulse Response Function (IRF): To transform discrete neural spikes into the typical morphology of a skin conductance response, we utilized the Bateman function. This mathematical model approximates the rise and decay of sweat gland activity. The function is defined as:

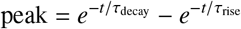

where t represents time, with rise time, (rise default = 0.7 s, during the rapid filling of the sweat ducts) and decay time (default = 3.0 s, corresponding to the slower reabsorption and evaporation process). The resulting IRF is normalized to a maximum amplitude of 1.0 to ensure consistent scaling across simulations.

Generating Synthetic Audience with Variable Individual Characteristics: To move beyond a single idealized trace, we instantiated a synthetic audience of 96 agents, matching the size of the real audience, by parameterizing individual agents with distinct physiological and attentional characteristics. For each agent, variability was introduced by sampling the following adjustable variation parameters from informed distributions: Each agent was assigned a base latency (*M*=*2*.*5s, SD*=*1*.*5s*) to account for neural transmission delays, and a responsivity factor (*M*=*1*.*0, SD*=*0*.*8*) to simulate varying levels of emotional reactivity. Furthermore, to model attentional lapses, we introduced a dropout rate. This parameter determines the probability that an agent fails to register a specific narrative event entirely, simulating moments of distraction. We also added a parameter that creates internal noise (e.g., task-unrelated thoughts), modeled as a Poisson process, and injected random spikes into the driver at a spont rate (0.05 to 0.3 Hz), with amplitudes drawn from an exponential distribution. Finally, a trial-to-trial jitter std (0.5 to 2.0s) was applied to event onsets to reflect the inherent stochasticity of biological timing. These characteristics served to make the model more realistic than a purely deterministic convolution, which would have created “identical clones”. Although these choices were based on current knowledge of audience response physiology, they represent preliminary settings rather than well-supported constants.

Neural Driver and Convolution: For each agent, a neural driver vector was constructed as a series of sparse impulses. These impulses consist of the AI-annotated movie events (weighted by their estimated intensity, see next section on how AI was used to generate these event-x-intensity series) and the agent’s unique spontaneous noise spikes. This driver was then convolved with the Bateman IRF to produce the raw phasic component of the GSR. Finally, each agent’s signal was z-scored to allow for population-level comparisons.

### Movie Stimulus Description and Content Analysis

This section introduces the example stimulus, Partly Cloudy (Sohn, 2009), and details our use of Multimodal AI Models to perform a psychological analysis and annotation of the stimulus. The animated short film Partly Cloudy (Sohn, 2009) is a 5-minute, 19-second dialogue-free narrative that follows Gus, a cloud who creates hazardous “offspring” (e.g., sharks, porcupines), and Peck, his stork delivery partner who incurs injuries while transporting them. The film was selected for its high-contrast emotional beats and its established history in research studying theory-of-mind and emotional processing (Richardson et al., 2018; Schmälzle et al., 2026).

We utilized AI to extract the psychological triggers, such as narrative intensity, physical threat, and emotional surprise, that serve as the elicitors for the simulated GSR events. Specifically, based on the rapid advances in AI for content annotation (e.g. Shen and Garg, 2025) and prior work in this area (Schmälzle et al., 2026), we reasoned that state-of-the-art multimodal models, such as Google’s Gemini, should be capable of “understanding” the movie (cf. Minsky, 2007). We thus crafted a prompt (see Supplementary Materials) and submitted the full video to Gemini Pro 3.1. Indeed, the output was meaningful, and the model was capable of extracting the most emotionally salient events (e.g. a high-arousal scene where a small crocodile almost bites off the protagonist’s head). Moreover, compared to previous experiments with multimodal AI models, we found that the timing of those events was also accurate. Table 1 contains a list of our movie-annotation along with a description of the scenes. Readers are advised to compare this against the video with the caveat that there are similar yet slightly different versions of the short film on the internet (https://www.youtube.com/watch?v=PfyJQEIsMt0).

**Table 1:** Description of AI’s Annotation for the Movie Events.

| Event # | Event Onset | Event Intensity | Scene Description / Justification |
| --- | --- | --- | --- |
| 1 | 10 | 1 | Scene fades in from black to the sky; visual orientation |
| 2 | 26 | 1 | Sudden lightning zap/audio pop as the kitten is created |
| 3 | 40 | 1 | Human baby suddenly starts crying loudly (auditory onset) |
| 4 | 55 | 1 | Camera reveals the dark cloud; music sharply shifts to ominous |
| 5 | 64 | 1 | Lightning zap creates the crocodile |
| 6 | 90 | 2 | Crocodile suddenly bites the stork’s beak (comedic violence/sharp visual) |
| 7 | 122 | 1 | Lightning zap creates the bighorn sheep |
| 8 | 124 | 2 | Bighorn sheep violently headbutts the stork (impact) |
| 9 | 160 | 1 | Lightning zap creates the porcupine |
| 10 | 163 | 2 | Porcupine lands on the stork; quills prick him (sharp reaction) |
| 11 | 192 | 2 | Shark is revealed; stork gasps and panics (strong emotional shift/fear response) |
| 12 | 218 | 3 | Massive thunderclap and lightning flash as the dark cloud erupts in crying (peak visual/audio/emotional arousal) |
| 13 | 244 | 1 | Stork unexpectedly returns into frame (orientation/relief) |
| 14 | 251 | 2 | Stork puts on the football helmet and pads (comedic beat) |
| 15 | 267 | 3 | Electric eel fiercely shocks the stork (bright flash, loud zap) |

### Performing the Simulation: Generating and Analyzing Synthetic Audience Data

Equipped with a detailed timeline annotation of salient movie events and the theoretical knowledge about EDA/GSR response generation, we proceeded to implement the simulation. A Jupyter-notebook to document the functions and reproduce all results, along with simulated and real GSR data is available at [https://github.com/nomcomm/generative/gsr]. In the following, we only summarize the key steps: First, we defined the timeline of significant events (see Table 1). Second, we specified the key function that would turn these events (which could be thought of as the activations of brain centers or the sudomotor nerve) into observable SCR responses. Specifically, this was done via a Bateman impulse response function. Next, this function was used to convolve the movie-event timeline to produce a timeseries of GSR data for one subject. Critically, because we didn’t only want to simulate one single subject, but an entire audience, we wrote an audience-generation function that included at its core the Bateman-IRF but also attempted to model other aspects (see details above). Our reasoning was that while the movie is basically the same stimulus for everyone (i.e., the same pixels and sounds arriving at the retina and the ear), we can expect individual differences due to various factors. These include variation in emotional reactivity, variability in sustained attention, differences in GSR-response speed or jitter, and so forth. These sources of individual variability were modeled to be generated from random or uniformly sampled distributions with parameters centered around estimates coming from the literature (e.g., Boucsein, 2013).

Variability was introduced by modulating the latency, responsivity, rate of spontaneous responses, response jitter, and dropout. The event-timeline was modified to include these factors and then convolved with the Bateman IRF to produce a GSR time series. With this basic response generation procedure, we then simulated GSR timeseries for an audience comprising 96 agents. See Schmälzle et al., 2026 for the code to reproduce this synthetic audience dataset.

### Existing Dataset of GSR Responses

To test the modeled GSR responses against GSR data collected from a read audience, we utilized a pre-existing dataset (Schmälzle et al., 2026) of electrodermal activity collected from human participants who viewed the same stimulus, the Pixar animated short film, Partly Cloudy (2009; 5 min 19 sec original version). The original dataset included 99 participants; however, three individuals were excluded from the final analysis due to data quality issues, resulting in a final sample of N = 96 (*Mage* = *19*.*91, SDage* = *2*.*05; 43 males*). This dataset serves as the benchmark, providing us with 96 time series of individual viewers’ GSR data, inter-subject correlation measures, as well as group-averaged GSR time series that allow us to identify the most evocative movie events. The GSR data were collected using iMotions software (Version 11.0.0.0; iMotions, 2025) with the Shimmer 3 GSR kit. Original data were sampled at 128 Hz and preprocessed (cleaned, aligned to the common movie onset/offset, resampled, and high-pass filtered at 0.05 Hz; see Schmälzle et al., 2026 for details).

## Results

### Generating a Synthetic Audience and Comparing Synthetic and Real GSR Traces

Figure 2 shows the main results. The top and middle panel display the individual data from the real (top panel) and synthetic (middle panel) as a carpet plot, meaning that the x-axis represents time and each line corresponds to one real/synthetic audience member, with color representing the amplitude of the preprocesses GSR signals.

**Figure 1.**
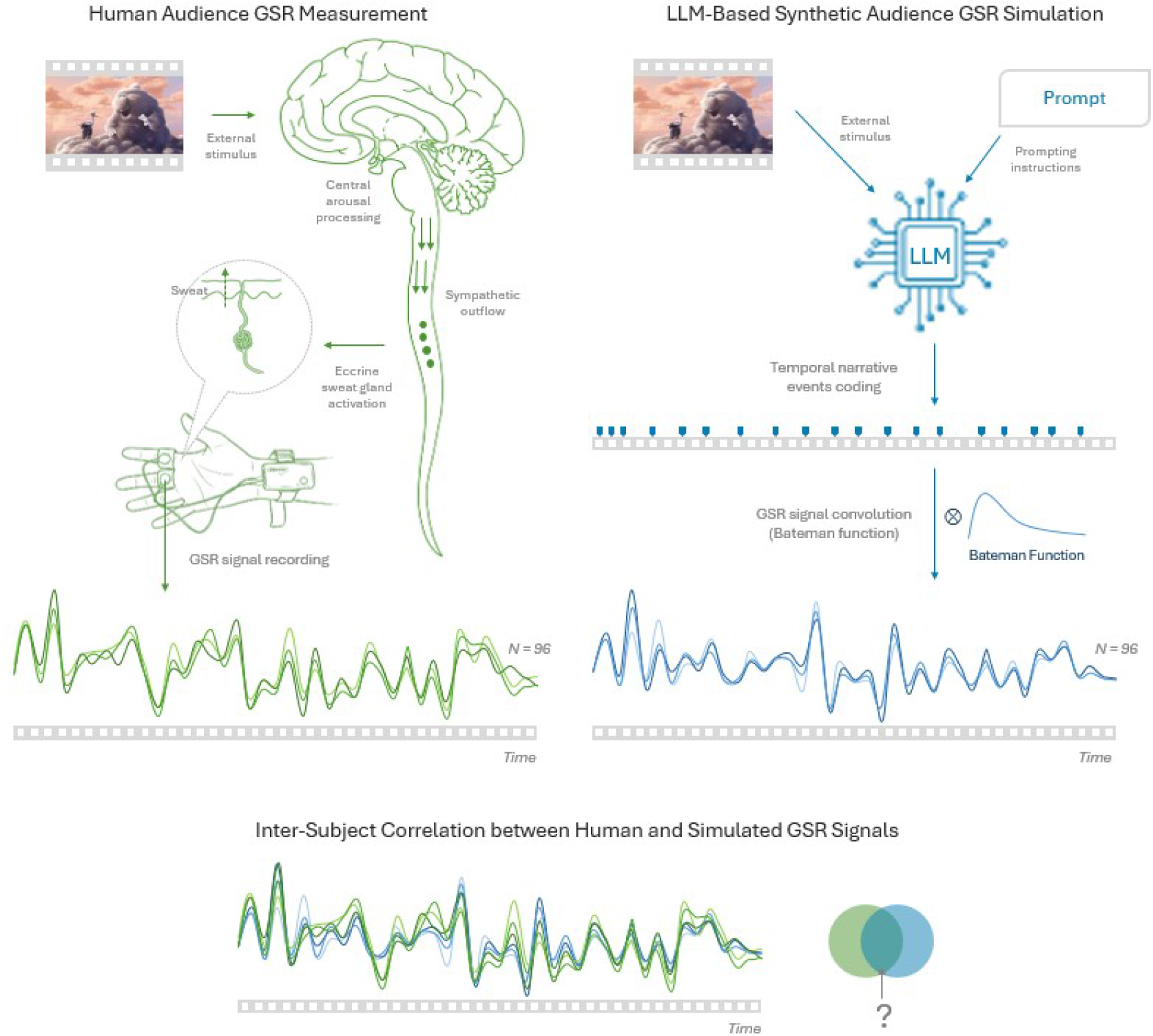
Conceptual Overview of the Idea. The figure illustrates the forward-modeling pipeline from dynamic narrative stimulus to simulated autonomic alignment. Temporally structured narrative events extracted from Partly Cloudy, https://www.youtube.com/watch?v=PfyJQEIsMt0, are represented as a continuous intensity time-series that serves as the latent neural driver of sympathetic activation. This driver is modeled as transient sudomotor bursts, which are convolved with a physiologically informed sweat gland impulse response function (Bateman function) to generate a continuous synthetic electrodermal signal. Multiple parameterized agents can be instantiated to simulate individual variability, producing a synthetic audience whose response similarity is quantified using inter-subject correlation (ISC). Simulated outputs are compared against empirical GSR data to evaluate model validity, estimate noise ceilings, and test mechanistic assumptions regarding stimulus–autonomic coupling.

**Figure 2.**
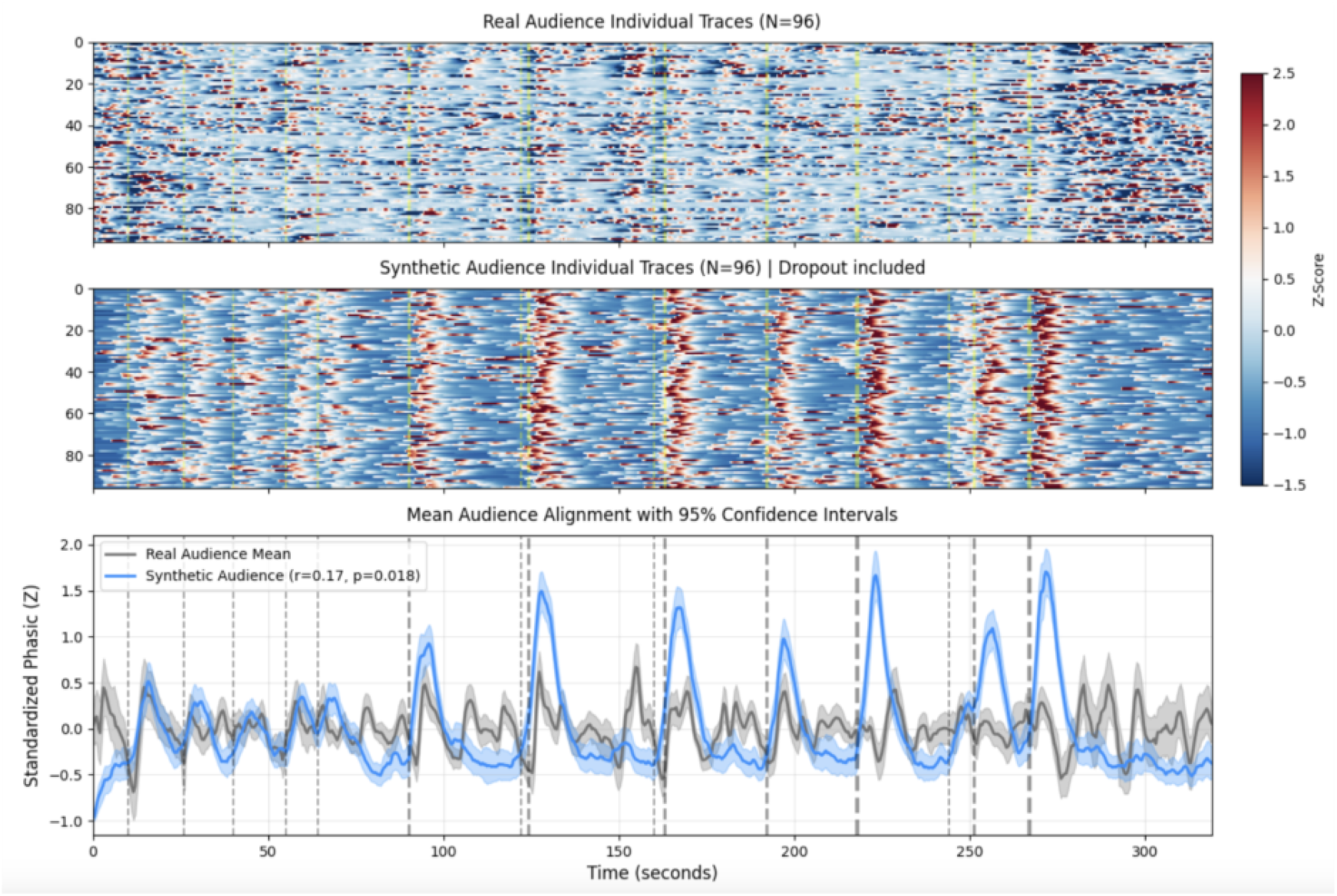
Empirical validation of the generative GSR model. Panel A displays the continuous synthetic GSR trace generated by the forward model alongside the empirical GSR signal across participants. Both signals are aligned to the temporal structure of the film, allowing visual comparison of event-locked phasic responses and slower tonic fluctuations. This shows the inter-subject correlation (ISC) time course derived from the synthetic audience and the real dataset. This panel evaluates whether the model reproduces the temporal fingerprint of autonomic alignment observed in human viewers. Peaks in ISC reflect moments of synchronized sympathetic activation across individuals. Results demonstrate the extent to which the forward-modeling architecture captures both phasic dynamics and higher-order collective synchronization patterns.

Turning to the simulated audience (middle panel), the depiction of the red-shaded strips (indicating GSR signal peaks) tend to align well with the ones in the real data, suggesting tentatively that our simulation captures something real. However, the inter-subject alignment of these peaks appears to be much higher. Indeed, when analyzing data from the synthetic audience, we find a higher ISC (see below).

The most critical result is depicted in the bottom panel of Figure 2 and refers to the direct comparison between the real and the simulated data. The line plots represent the group-averages (and variability) of the data depicted above, i.e. creating the group-averaged GSR signal for both the real and simulated audiences, respectively. As can be seen, several salient peaks are expressed both in the real datasets and –more notably due to the higher inter-subject alignment – in the synthetic audience. Statistically, this similarity in the real and simulated time-series is supported by a correlation of *r* = .*17*. This correlation is obviously not perfect, nonetheless it is significant.

### Simulating Many Audiences to Show that Synthetic-vs.-Real GSR Alignment is Robust

We note, however, that while our approach to simulate a synthetic audience of 96 participants is already quite realistic, it is of course the case that – as with all simulations – a new run of the simulation would yield slightly different results. Thus, the number provided above, a correlation of *r* = .*17* between the simulated and real data, is itself coming from a distribution. While our statistical test did not only correlate these two traces, but we performed a more sophisticated assessment of significance against shuffled/time-shifted surrogate data, we still wanted to conduct a more comprehensive analysis. To that end, we re-ran the entire audience simulation procedure (i.e. generating a N = 96 audience) 1000 times. The result of this simulation is shown in Figure 3, demonstrating that our correlation lies in the middle range of this distribution, which ranges from about .1 to about .2 at maximum. Importantly, these permutations were done with the same parameter configurations mentioned above (see Methods; i.e. the fixed parameters for time-delay, emotional reactivity, etc.). We did not conduct further parameter sweeps of these hyperparameters, but this is obviously an area for future research.

**Figure 3.**
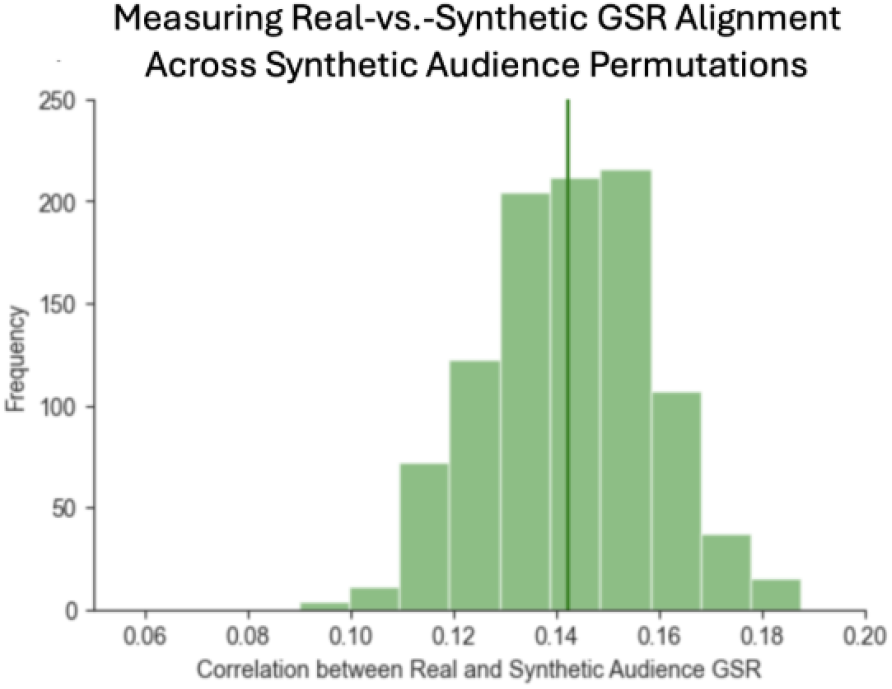
Using Resampling to Generate 1000 Synthetic Audiences. Shown is a histogram of the resulting correlations between the synthetic audience and the real audience data. A total of 1000 synthetic 96-person audiences were generated, and the resulting correlations with the real audience were computed.

### Inter-Subject Correlation Analysis of Synthetic and Real GSR Data

As can be seen in the top panel of Figure 2, there is a clear alignment of the GSR traces across the 96 real-audience datasets; and looking at the middle panel, which shows the synthetic audience data, the alignment appears even more clearly. To quantify this alignment, we computed inter-subject correlation analysis. A split-half approach was utilized for visualization, meaning that we randomly split the group of 96 viewers into two halves, averaged the time-series across viewers, and plotted both resulting group-averaged time series. The result of this analysis are shown in Figure 4 (blue traces for real audience, red for synthetic audience). As can be seen, the alignment for the real audience was robust (about .5-.6 depending on the specific split), but the synthetic audience alignment was even higher (>.9). One challenge of this split-half visualization is that the specific way an audience of 96 is split into two halves (e.g. 1:48, 48:96) is somewhat arbitrary. To overcome this and get a clearer view of the variability and sampling distributions of the real and synthetic data, we also permuted this split-half-ISC analysis 1000 times. The resulting distributions as well as the grand means are shown in Figure 4. As can be seen, the real data split half ISC lies around .63, whereas the split half ISC for the synthetic audience lies at >.9. Clearly, the distributions do not even overlap, suggesting that the synthetic audience is more aligned within itself than the real data, which are more variable. This suggests that while our simulation is capturing something real about the underlying GSR data (see analyses above demonstrating the correlation between real and synthetic data), the synthetic data are still too similar to each other – pointing to unmodeled sources of error (measurement error, internal variance, variability between subjects).

**Figure 4.**
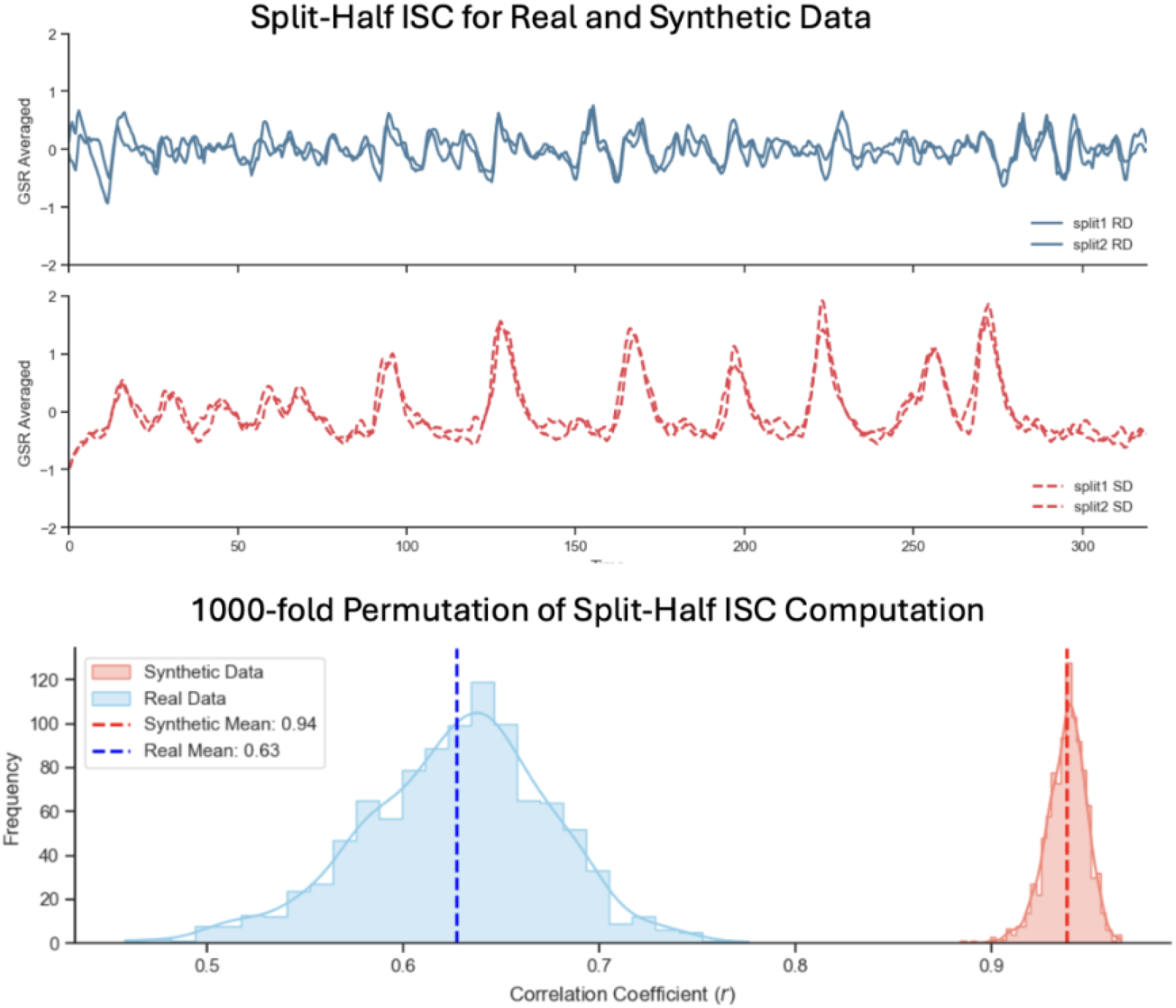
Split-Half ISC Analysis Results. Top panels (blue and red): Time series plot for one split-half (viewers 1-48 vs. viewers 49:96) for both real and synthetic audiences, revealing correlations of around .6 and .95. Bottom panel: Plot of the distributions arising from resampling the split-half procedure, along with the resulting grand means.

## Discussion

The results of this study are clear and can be summarized as follows: First, two different AI models (Gemini Pro 3.1 and initially ChatGPT-5.2 for proof of concept) were exceedingly capable of provided a finely annotated and time-stamped list of relevant movie events. This in itself marks progress compared to just one year ago where precursor models were not yet able to “understand” see Minsky, 2007 for a caveat regarding the word “understand” in AI modeling, particularly the social-cognitive content of dynamic stimuli. Only a few years ago, computervision pioneer Karpathy used a similar task to claim that “we are really far away” from AI vision models that could competently analyze social-cognitive image content (Karpathy, 2012); however, at this point, AI’s capabilities have exponentially advanced to make this feasible.

Perhaps the most important finding relates to the correspondence between real and simulated audience GSR responses. The convolution of the AI-annotated events with a physiologically plausible Bateman IRF produced results that were significantly similar to the real GSR time series. This is our main finding and, to our knowledge, is the first time that such a computational approach has been applied to GSR in media psychophysiology. Prior work has largely been focused on modeling discrete events like SCRs in conditioning paradigms (e.g. Bach, 2014b; Benedek and Kaernbach, 2010b), but our work here shows that when combined with the capabilities of modern AI for content analysis, this approach can now be expanded towards narrative media, drastically expanding the scope and application potential of this enterprise.

While the overall goal to plausibly model GSR data was achieved by showing that they correspond with real data, it is important to note that this correspondence was still not perfect: Despite our efforts to introduce further biological plausibility by modeling e.g. audience reliability, response speed, and attentiveness, Figure 2 clearly shows two key differences between real and modeled data: First, comparison of the carpet plots shows that the simulated audience is far more homogenous in itself than the real audience. Thus, although we modeled individual variability, the real audiences’ variability was still greater. Second, inspection of the bottom panel in Figure 2 shows that while our model seems to capture the largest peaks (e.g. the crocodile-bites-off –head scene and similar others), there are still notable discrepancies in the timing and magnitude of smaller fluctuations, suggesting that more subtle, transient features of the stimulus are remain to be fully accounted for.

## Theoretical Implications

The implementation of a generative model for GSR addresses a critical need for theoretical precision within the communication field regarding social-affective responses to dynamic stimuli (DeAndrea and Holbert, 2017). This provides a bridge between the realm of media psychophysiological research, computational communication, and agent-based modeling (Smaldino, 2023; Carpenter et al., 2024), introducing the new concept of a synthetic, simulated audience.

As emphasized by DeAndrea and Holbert, 2017, theoretical progress is often stalled by a lack of precise terminology and lack of explicitness in specifying the “how” of theoretical entities interact to produce observed effects. With this in mind, this generative approach provides a common language that translates narrative events into specific biophysical parameters. Instead of treating the relationship between a media message and its physiological effect as an implicit black box, the model necessitates an explicit account of the underlying mechanisms, from the discrete firing of the sudomotor nerve to the physical constants of the sweat gland impulse response. By transitioning from descriptive and umbrella-term labels like arousal toward an increasingly explicit computational framework, the simulation code itself functions as a formal model of the psychophysiological responses within the message reception process.

Utilizing a forward-modeling architecture, we move beyond simple correlation to test the boundaries of communication theory in a controlled, synthetic environment. This supports a mechanism-based theorization, as the model allows for the formal definition of the neural driver as the active interface between a structured message and the receiver’s body. Moreover, by parameterizing spontaneous sudomotor activity and reaction jitter, the model provides a framework for understanding why autonomic alignment is naturally lower than direct sensory measures like gaze (e.g. due to Receiver Noise, Physiological Constraints, and Measurement Noise). Finally, this furthers theoretical advancement indicated by the shift in media psychophysiology from a reactive discipline to a predictive one, allowing for the simulation of what-if scenarios regarding receiver states or message structures potentially across a variety of mediums including live action film, animation, music, educational content, or even in the instance of temporal rescaling (Pearl and Mackenzie, 2018).

### Practical Implications

One of the most immediate practical benefits of this generative approach is the ability to create high-fidelity synthetic datasets with the potential to predict real audience responses and subsequent behavioral outcomes. More so, researchers can use these simulations to test the sensitivity and robustness of their analysis pipelines before ever recruiting human participants. By generating synthetic audiences with known levels of responsivity and spontaneous noise, researchers can gauge the minimum effect size required to reach statistical significance in audience studies and simulate how audiences would respond to various manipulations.

The possibility of generating synthetic audiences basically represents an extension of the Agent-Based Modeling framework to the field of audience behavior and audience reactions (cf. Smaldino, 2023; Wilensky and Rand, 2015; Carpenter et al., 2024). Specifically, the ability to parameterize the neural driver could allow this framework to be integrated into ABM, leading to more realistic simulations (Epstein, 2014). In doing so, it reorients the analysis from abstract, macro-level assumptions toward audience dynamics that are directly grounded in physiologically validated data.

Indeed, one critique of ABM is that they often abstract away from implementational detail, treating nodes in a network as simplified entities rather than “real humans”. In this book “Agent Zero”, modeling pioneer Epstein, 2014 therefore called for more research on the neurocognitive foundations of agents and simulations; our work here provides exactly this missing bridge between the individual level of bodily reactions and the collective aggregate of audience response dynamics. Thus, rather than treating an audience as a monolithic aggregate, we can simulate thousands of virtual receivers with varying physiological profiles (e.g., reactive vs. distracted). This could allow for the study of how specific narrative structures might propagate through different audience segments. This approach has the potential to allow for the parameterization of audience segments along different dimensions, enabling the examination of how narrative structures differentially engage viewers with distinct cognitive and emotional profiles.

In the long run, the GSR modeling framework might even be further extended to incorporate culturally or developmentally informed response priors, capturing how socially learned norms and developmental effects shape media perception and response. Such extensions provide a pathway for modeling how shared physiological responses emerge within groups while systematically varying between segments of the population to comprehensively evaluate individual differences in audiences.

This framework also effectively transforms simulated audience responses into a continuous annotation of the media content. By linking specific narrative features identified by LMMs, to the resulting physiological traces, media creators can perform a functional audit of their work without the need for message testing with live audiences. This has significant potential for content optimization, such as the identification of dead spots or moments where the collective grip of the message begins to fail, and ISC starts to diverge. In applied contexts, this approach may be particularly useful for short-form media currently seen dominating social media today, where limited temporal windows place a premium on fine-tuning the placement of high-impact moments to maximize audience retention, emotional impact, and sustained audience-alignment without rapid divergence in engagement.

### Strengths and Limitations

The primary strength of this work lies in the computational precision it brings to media psychophysiology, enabling the formalization of dynamic autonomic responses as a generative process rather than a purely descriptive signal. Furthermore, by validating the simulation against a large-scale, real-world dataset, we demonstrate the model’s predictive validity and its capacity to reproduce the complex temporal dynamics observed in human audiences, providing a scalable framework for linking stimulus structure to shared physiological responses.

Despite its generative power, the model remains a simulation and possesses several inherent limitations. The current framework assumes a fixed impulse response function (the Bateman function) across all subjects. In reality, the shape of a sweat burst can be influenced by individual biological factors such as skin temperature, hydration, and varying ductal filling rates, just like the BOLD signal does. Thus, this simplification enables tractable modeling but necessarily abstracts meaningful interindividual differences in electrodermal dynamics. A related limitation concerns the modeling of the tonic component. While the random walk simulates the tonic component of GSR, it does not currently account for how slow-moving baseline shifts might be systematically driven by global narrative states, such as sustained suspense or relief, rather than purely stochastic drift. This is perhaps one of the biggest limitations at this point, but we wanted to first establish the validity of SCR prediction before moving on to more complicated fluctuations over longer timescales with skin conductance levels.

Future research should expand this generative logic to other modalities of the autonomic system to create a more holistic synthetic receiver/audience. Integrating these generative physiological profiles into agent-based models could allow researchers to simulate the impact of media messages on large, diverse populations with varying levels of responsivity and internal noise. Next, as Large Multimodal Models (LMMs) and Machine Learning more broadly continue to evolve, they can be used to perform more nuanced, frame-by-frame psychological reverse-engineering of content, providing a more detailed design matrix to drive the physiological simulation. Additionally, the current framework does not explicitly incorporate continuous stimulus features, such as luminance, contrast, or motion energy, as well as discrete structural events like sudden cuts and other physical surprises. These features can be conceptualized as low-level forms of perceptual saliency, suggesting a natural extension of the model toward integrating both affective and sensory drivers of autonomic dynamics.

### Summary and Conclusions

This study demonstrates that the audience’s physiological reaction to a message is not merely a collection of idiosyncratic responses, but a structured alignment induced by the narrative signal itself. By implementing a generative, forward-modeling framework, we have shown that it is possible to move beyond descriptive analysis to a mechanistic account of how narrative beats drive sudomotor activity over time. More broadly, this approach bridges computational modeling, media psychology, and social cognition, offering a scalable framework for studying collective experience as it unfolds in real-world media environments.

## Data

The anonymized dataset and data analysis scripts are available via GitHub at https://github.com/nomcomm/generative_*g*_ *sr*

## Ethics Approval

The study was approved with the IRB of Michigan State University.

## Consent to Participants

All participants provided written informed consent.

## Use of Generative AI

Google Gemini and Nano Banana were used during the drafting process to assist with the organization of the manuscript, the refinement of theoretical arguments, copy-editing, and creating elements of the figures. AI was also used to optimize code. All original ideas, hypotheses, data analysis, and final theoretical conclusions remain in the intellectual property of the authors. The authors reviewed and edited all text and code and take full responsibility for the content.

## Supplementary Materials

The following prompt was used with Gemini Pro 3.1 to generate the timeline of movie events used for the model.

Role: You are an expert in psychophysiology and affective computing.

Task: Analyze the attached 5-minute video stimulus to identify discrete “arousal events” that would trigger a Phasic Skin Conductance Response (SCR).

Coding Criteria:

Identify Onset: Note the exact second (timestamp) a stimulus begins that would cause a startle, an orientation response, or a sharp emotional shift (e.g., a jump scare, a loud noise, a sudden visual change, or a highly emotional dialogue beat).

Assign Intensity: On a scale of 0.5 to 5.0, estimate the “Neural Strength” of the stimulus.

0.5 - 1.5: Minor orientation (a new character enters, a phone rings).

1.6 - 3.5: Moderate arousal (an argument starts, a tense moment).

3.6 - 5.0: Peak arousal (a physical shock, a major plot twist, a loud explosion).

Output Format: Please provide the data strictly as a Mark-down Table and then as a JSON list so I can parse it into my Python/MATLAB model.

JSON Structure Example: [“timestamp sec”: 92, “intensity”: 2.0, “description”: “brief description of event”]

Constraints:

Do not include the 1 to 5 second physiological lag; I want the stimulus onset (the “neural” event). My forward model will handle the convolution and lag.

Ignore the tonic (slow-moving) background; focus only on discrete phasic triggers.

